# Arginine methylation of Puf4 drives diverse protein functions

**DOI:** 10.1101/2022.06.21.497104

**Authors:** Murat C. Kalem, Sean Duffy, Shichen Shen, Jan Naseer Kaur, Jun Qu, John C. Panepinto

## Abstract

The evolutionarily conserved Pumilio domain-containing RNA binding proteins (RBPs) are involved in many steps of post-transcriptional gene regulation, including RNA stability, polyadenylation, deadenylation, and translation. RBPs are post-translationally modified by methylation of arginine/glycine-rich domains, though the consequences of these modifications are not well known. We determined the arginine methylation and phosphorylation landscape of the Pumilio domain-containing RBP Puf4 from the basidiomycete fungus *Cryptococcus neoformans*. We found that methyl-deficient Puf4 mutants do not complement critical *PUF4* deletion phenotypes, such as resistances to endoplasmic reticulum stress and antifungals, and cell wall remodeling. Methyl-deficient mutants also exhibit unique RNA and protein interactions. Lastly, we identified intra-protein cross talk between post-translationally modified methylated and phosphorylated residues. Overall, we show that post-translational modifications, particularly arginine methylation, of Puf4 regulate the functions of this RBP.

## Introduction

RNA binding proteins (RBPs) are key post-transcriptional regulators that fine-tune critical cellular events and are instrumental in every step of the RNA life cycle, from synthesis to translation and decay. Misregulation of RBP activity can be pathological, contributing to cancer and neurodegeneration ^1–3^. The binding domains within RBPs and the cognate RNA features they recognize are evolutionarily conserved across kingdoms, but the processes they regulate and their consequences may be kingdom specific ^4,5^. Pumilio and Fem-3 mRNA binding factor (PUF) domains in RBPs, first identified in the arthropod *Drosophila melanogaster* and the nematode *Caenorhabditis elegans*^6–8^, contain highly conserved helical repeats that regulate the localization, decay, and translation of specific mRNAs ^9,10^. *D. melanogaster* has one PUF domain-containing RBP, humans have two, the basidiomycete fungus *Cryptococcus neoformans* has four, the ascomycete fungus *Saccharomyces cerevisiae* has six, and the nematode *C. elegans* and the kinetoplastid parasite *Trypanosoma cruzi* each have ten ^9–12^. These different numbers of PUF proteins suggest evolutionary diversification in the regulatory roles these proteins play ^13,14^.

RBPs are dynamically regulated by post-translational modifications (PTM) ^15–17^. For example, the highly conserved and abundant arginine-glycine (RG)-rich domains in RBPs can be methylated ^18,19^. Arginine methylation is catalyzed by arginine methyltransferases (RMTs), which are similarly diverse across the evolutionary landscape ^20,21^. The *S. cerevisiae* genome encodes four RMTs, whereas the human genome encodes nine, contrary to their numbers of PUF proteins (six and two, respectively) ^22^. The apparent inverse association between the numbers of RMTs and PUF proteins led us to hypothesize that functional diversification of gene regulation is driven by the variety of RBPs in less complex organisms and by arginine methylation (and PTMs in general) of RBPs in more complex organisms. The ability to leverage dynamic and often reversible PTMs may impart agility for post-transcriptional gene regulation ^23,24^. Our recent work to characterize RMTs in *C. neoformans* revealed that one of its four PUF proteins, Puf4, interacts with Rmt5, suggesting that its function is controlled through methylation ^25^. Puf4 regulates mRNAs involved in cell wall biosynthesis and is involved in the splicing of *HXL1,* the major endoplasmic reticulum (ER) stress transcription factor ^26,27^. Of note, *puf4*Δ mutants are resistant to the ER stress-inducing drug tunicamycin and the antifungal drug caspofungin ^26–28^.

Although PUF proteins have been extensively studied, we are not aware of any studies on the regulation of PUF protein function by PTMs ^9,30,32^ The human PUF protein Pum1 was shown to be methylated, but the functional importance of this modification was not explored ^29^. In this study, we set out to assess Puf4 PTMs in *C. neoformans* and their functional consequences. *C. neoformans* is a well-established and genetically tractable fungal model organism that has been used to investigate post-transcriptional processes and to showcase their divergence across fungal clades ^31,33,34^.

## Results

### Evolutionary divergence of RG domains and arginine methylation in *C. neoformans* Puf4

To investigate the evolutionary conservation of critical domains in Puf4 across the fungal kingdom, we included fungi representing different taxonomic divisions. A phylogenetic analysis revealed that the PUF domain is highly conserved in fungi, but disordered domains and RG-rich domains are more divergent (Figure 1a). This suggests a divergence in Puf4 function, because disordered domains evolve faster and are important for liquid–liquid phase separation ^15,35^. The RG-rich domains appeared to be unique to each species (Figure 1b, highlighted by the blue box) and thus could either be post-translationally modifiable domains that are not under constant selection or have species-specific roles.

**Figure 1.**
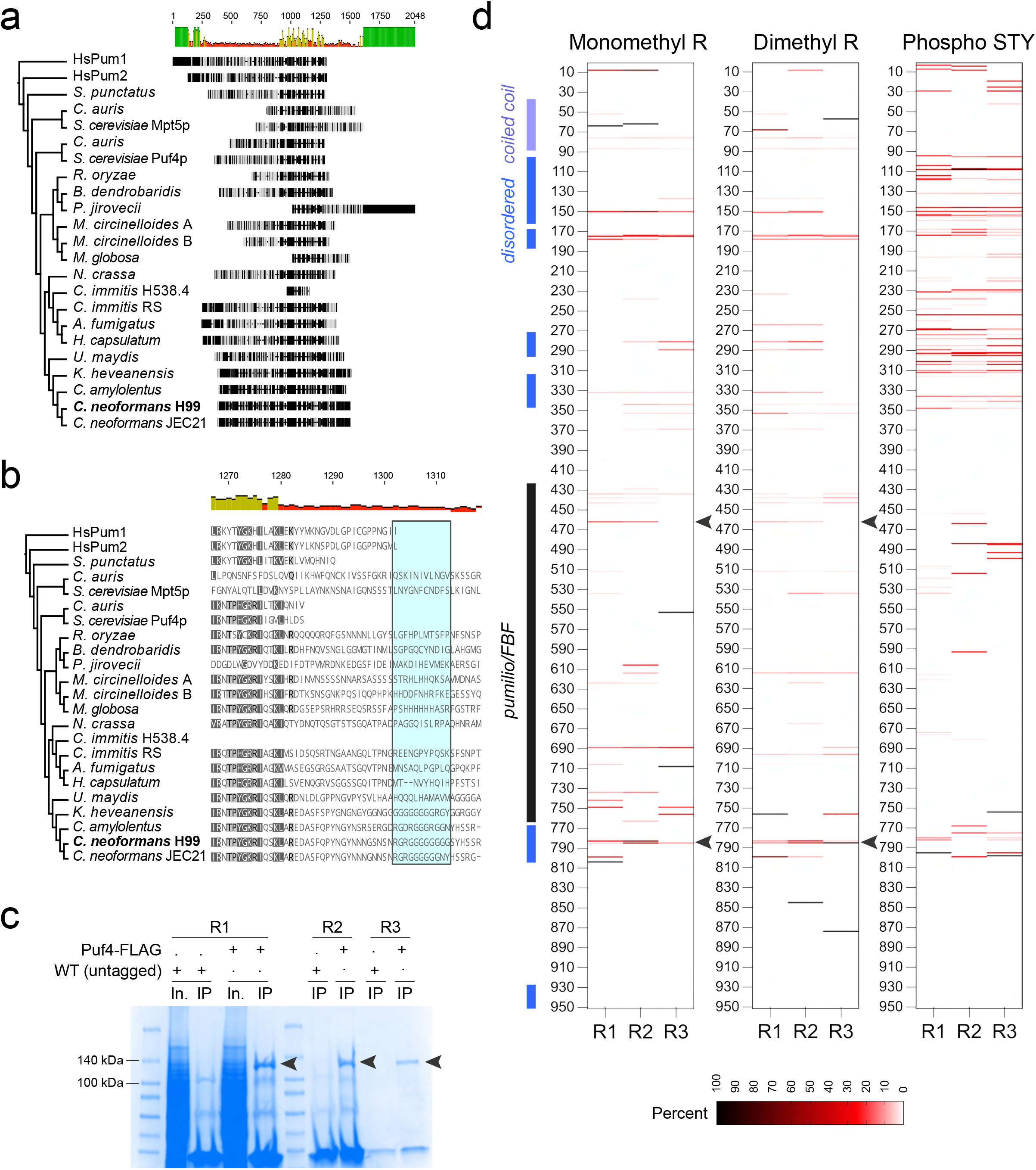
Evolutionary divergence of RG domains and arginine methylation in *C. neoformans* Puf4. (a) Phylogenetic analysis of Puf4 across fungi. Protein sequences used for analysis were exported from NCBI (compiled in Table S1), and phylogenetic analysis was performed using Geneious. (b) Evolutionary divergence of RG-rich domains. The blue box shows the RG-rich putative methylation hub C terminal to the PUF domain. (c) Immunoprecipitation of Puf4. Coomassie-stained gel shows three replicates of immunoprecipitated Puf4-FLAG. Arrowheads show the Puf4-FLAG bands that were excised and analyzed for PTMs using label-free LC-MS/MS. (d) PTM landscape of Puf4. Heat maps show the percentage of modifications at each amino acid site for monomethyl arginine methylation, dimethyl arginine methylation, and STY phosphorylation. Percentage of modifications for each amino acid was calculated by dividing the number of modifications at one site by the number of times that site was detected among all peptides. Important protein domains are noted on the left along with the amino acid numbers.

To characterize the PTM landscape of Puf4, we performed label-free liquid chromatography-tandem mass spectrometry (LC-MS/MS) of Puf4 immunoprecipitated from lysates of a *C. neoformans* strain overexpressing Puf4. Specifically, we mapped the monomethyl and dimethyl arginine modifications across the protein sequence (Figure 1c, bands excised from the gel are indicated with arrowheads). Because Puf4 is downstream in the calcineurin pathway and is dephosphorylated by the phosphatase Cna1^36^, we also mapped the phosphorylation of serine/threonine/tyrosine (STY) residues. This approach yielded 81.05%, 74.50%, and 74.43% Puf4 sequence coverage in three biological replicates. The percentage of modifications per amino acid site was calculated by dividing the number of times a site was modified by the total number of times the same site was detected across sampled tryptic peptides. The results revealed that *C. neoformans* Puf4 is methylated and phosphorylated under basal conditions (Figure 1d). The methyl marks were distributed throughout the protein, but there was strong methylation of arginines in the disordered domains, with high confidence across the three biological replicates. The species-specific RG-rich domain C terminal to the PUF domain (i.e., the methylation hub) that we identified in our phylogenetic analyses exhibited both monomethyl and dimethyl modifications. The proximity of this methylation hub to the RBD and its presence within a disordered domain suggest that arginine modification regulates protein function. By contrast, very few of the arginine residues in the PUF domain were methylated, suggesting that these sites are important for critical protein–protein interactions. The phosphorylation of STY residues was similarly concentrated in the disordered domains (Figure 1d). Thus, both arginine methylation and STY phosphorylation in Puf4 likely influence proteinprotein or protein–RNA interactions.

### Mutations for methyl deficiency of critical residues impact the complementation of Puf4 deletion phenotypes

We picked two highly modified residues to further probe the PTM–function relationship: Arg783/Arg785, located within the RG-rich methylation hub C-terminally adjacent to the PUF domain (Figure 2a), and Arg462, located within the PUF domain (Figure 2c). We mutated these arginines to lysines (RtoK) to establish cell lines expressing methyl-deficient Puf4. We also used FLAG-tagged wild-type and methyl-deficient Puf4 strains to complement the Puf4 deletion (*puf4*Δ) strain.

**Figure 2.**
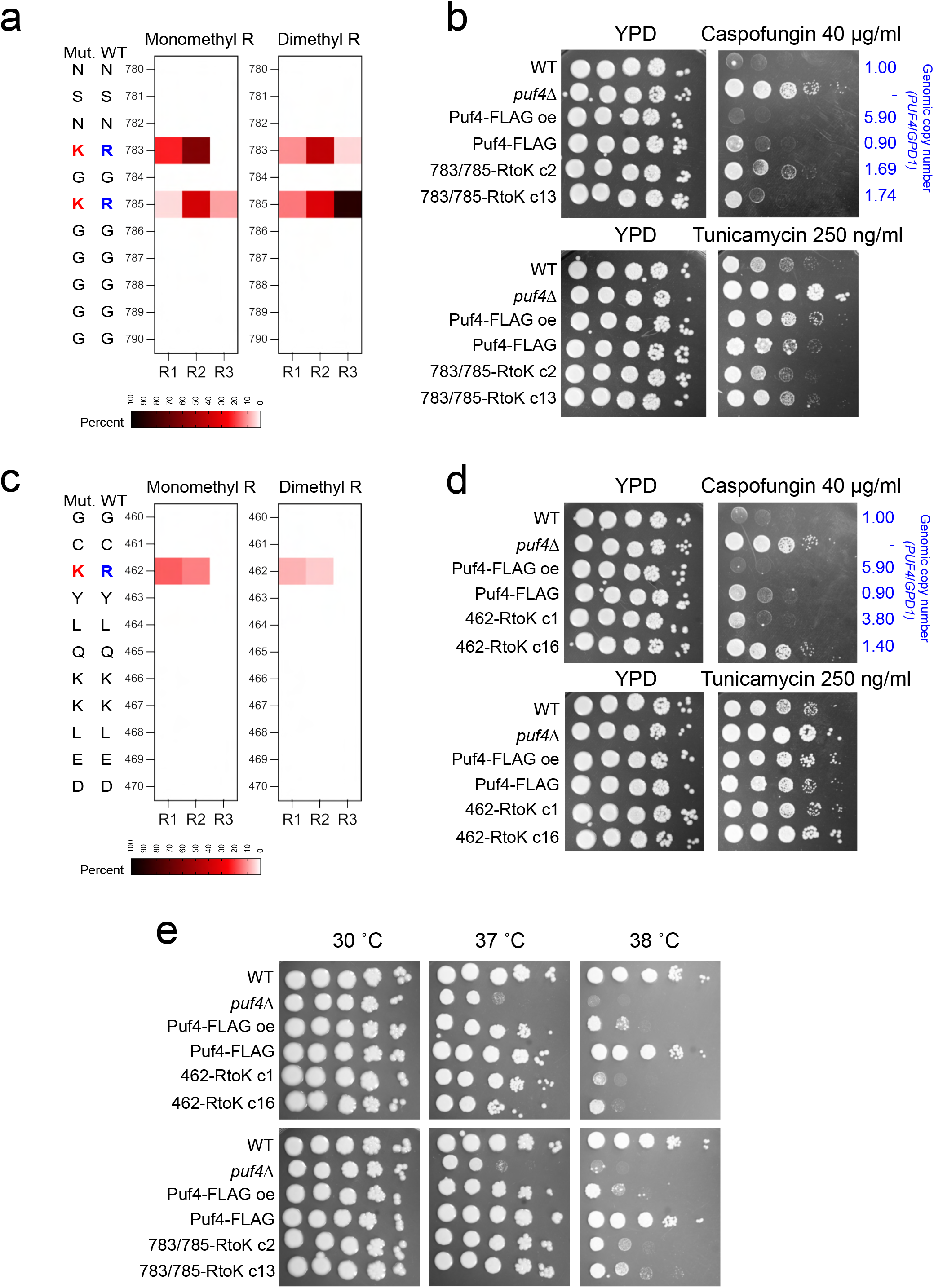
Mutations for methyl deficiency of critical residues impact the complementation of Puf4 deletion phenotypes. (a) Arginine methylation of the RG-rich domain at the N-terminal end of the RNA-binding domain. Zoomed-in map shows the level of methylation of arginines at sites Arg783 and Arg785. (b) Complementation of the *puf4*Δ mutant with the Puf4-FLAG 783/785 RtoK methyl-deficient mutant. Cells were analyzed for complementation of caspofungin and tunicamycin resistance phenotypes. The genomic copy number was determined by RT-qPCR using genomic DNA as a template to estimate the number of complementation constructs integrated. *GPD1* was used as the normalization gene. (c) Arginine methylation of the PUF domain at Arg462. (d) Complementation of the *puf4*Δ mutant with the Puf4-FLAG 462 RtoK methyl-deficient mutant. Cells were analyzed for complementation of caspofungin and tunicamycin resistance phenotypes. (e) Spot plate dilution assays at 30°C, 37°C, and 38°C.

We established two independent complement strains for each mutation and investigated previously reported caspofungin and tunicamycin resistance phenotypes ^26,27^. We accounted for the integration copy number of ectopically expressed wild-type or mutant Puf4-FLAG constructs, which can bias the interpretation of results. Both clones (c2 and c13; with similar copy numbers) of the Puf4-FLAG 783/785 RtoK strain partially complemented the caspofungin resistance phenotype and fully complemented the tunicamycin resistance phenotype (Figure 2b). By contrast, only one of the Puf4-FLAG 462 RtoK clones (c1) complemented both caspofungin and tunicamycin resistance phenotypes (Figure 2d); however, the integration copy number analysis showed that clone 1 is an overexpression strain with four integrations, whereas clone 16 likely carries a single genomic integration of the mutant construct. Thus, overexpression of Puf4-FLAG 462 RtoK complemented the deletion phenotype, but a single copy integration was not sufficient. We conclude that the Puf4-FLAG 462 RtoK methyl-deficient protein is only partially functional, which can be surmounted with increased dosage.

Because *PUF4* deletion causes a defect in thermotolerance of *C. neoformans,* which is critical for the pathogenicity of the fungus, we investigated the effect of methylated residues on growth at different temperatures. Both methyl-deficient Arg462 and Arg783/Arg785 mutants complemented the thermotolerance defect in the *puf4*Δ strain at 37°C but not at 38°C (Figure 2e). This suggests that modifications of Puf4 may have temperature-dependent functions at 37°C and 38°C. Overall, our results highlight that the identified methylated arginine residues modulate RBP function through yet unknown mechanisms under compound and temperature stresses.

### Mutation of Arg462 for methyl deficiency abolishes the ability of Puf4 to modulate RNA stability and cell wall maintenance

Puf4 binds to cell wall biosynthesis mRNAs and stabilizes these transcripts ^26^. We investigated if methyl-deficient Puf4 stabilized two key cell wall biosynthesis transcripts: *FKS1,* encoding ß-1,3-glucan synthase, and *CHS4,* encoding chitin synthase 4 ^37^. We investigated the stability of *FKS1* and *CHS4* under basal conditions at 30°C following transcription shutoff. Both single-copy and overexpression Puf4-FLAG constructs complemented the RNA decay phenotype of the deletion strain (Figure 3a; see also Figure S1). Clone 1 but not clone 16 of the Puf4-FLAG 462 RtoK mutant also complemented the mRNA decay phenotype, likely because of its higher gene dose. Both clones of Puf4-FLAG 783/785 RtoK complemented the mRNA decay phenotype.

**Figure 3.**
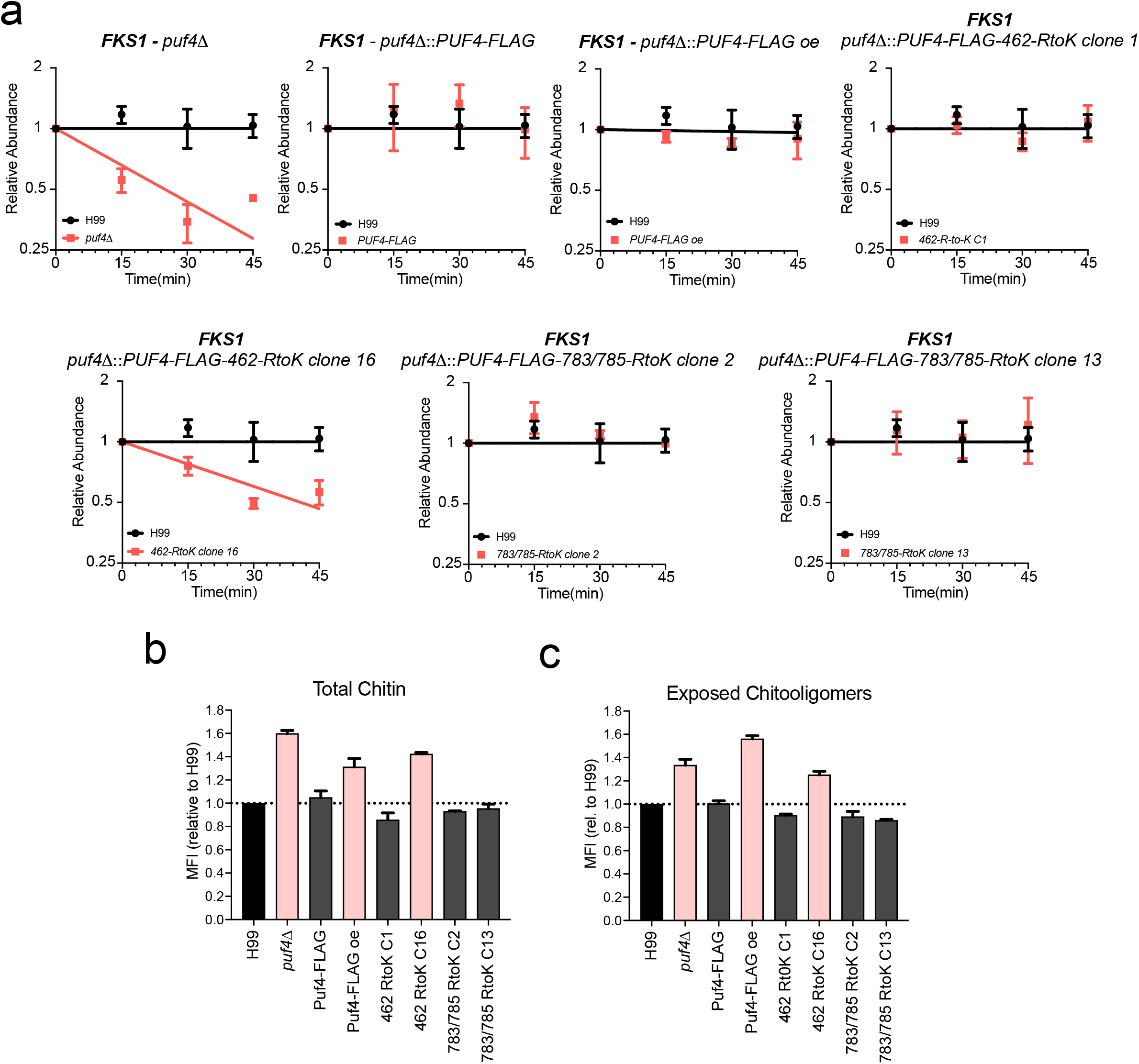
Mutation of Arg462 for methyl deficiency abolishes the ability of Puf4 to modulate RNA stability and cell wall maintenance. (a) *FKS1* mRNA stability. *FKS1* transcript abundance was determined by RT-qPCR following transcription shutoff via 1,10-phenanthroline. *GPD1* was utilized as the control gene for normalization. Means from three replicates were plotted, and differences between strains were analyzed using one-phase exponential decay analysis. See also Figure S1. (b,c) Cell wall chitin levels. Cells were grown to the mid-log stage and stained with calcofluor white and wheat germ agglutinin. Fluorescence was detected via flow cytometry, and mean fluorescence intensities (MF) are plotted.

We previously found that cell wall chitin and exposed chitooligomer levels are increased in the *puf4*Δ strain ^26^, and similar assessments revealed that this phenotype was complemented by a single copy of wild-type Puf4-FLAG but not by an overexpression construct (Figure 3b and 3c). This suggests that Puf4 dosage is critical for some phenotypes. In addition, both Puf4-FLAG 783/785 RtoK mutant clones (but not the Puf4-FLAG 462 RtoK mutant) complemented the chitin and chitooligomer phenotypes. Overall, these data suggest that mutation of Arg462, but not Arg783/785, abolishes the Puf4 function related to cell wall maintenance in a gene dosedependent manner.

### Diverse modulation of gene expression and RNA binding by Puf4 in *C. neoformans*

We utilized a transcriptome-wide approach to investigate the global role of Puf4 in gene expression regulation. RNA-seq analysis of *puf*4Δ cells grown to mid-log phase at 30°C revealed 585 up- and 633 downregulated transcripts (relative to expression in the wild type) (Figure 4a). Interestingly, these changes included 7 up- and 112 downregulated long noncoding RNAs (lncRNAs), suggesting that Puf4 drives the expression of a small population of lncRNAs. We utilized gene ontology (GO) analysis to investigate the biological processes, cellular components, and molecular functions that are represented in the Puf4 regulon in *C. neoformans*. Analysis of upregulated transcripts included GO categories such as transcription regulator complex, nucleus, ribonucleoprotein complex biogenesis, ribosome biogenesis, and RNA metabolic process (Figure 4b). GO categories for the downregulated transcripts included the intrinsic component of membrane, carbohydrate metabolic process, transmembrane transport, and hydrolase and oxidoreductase activity (Figure 4c). Thus, Puf4 appears to suppress the expression of genes that belong to post-transcriptional and transcriptional regulation-related categories and enhance that of genes related to membrane components, carbohydrate metabolism, and protein activity.

**Figure 4.**
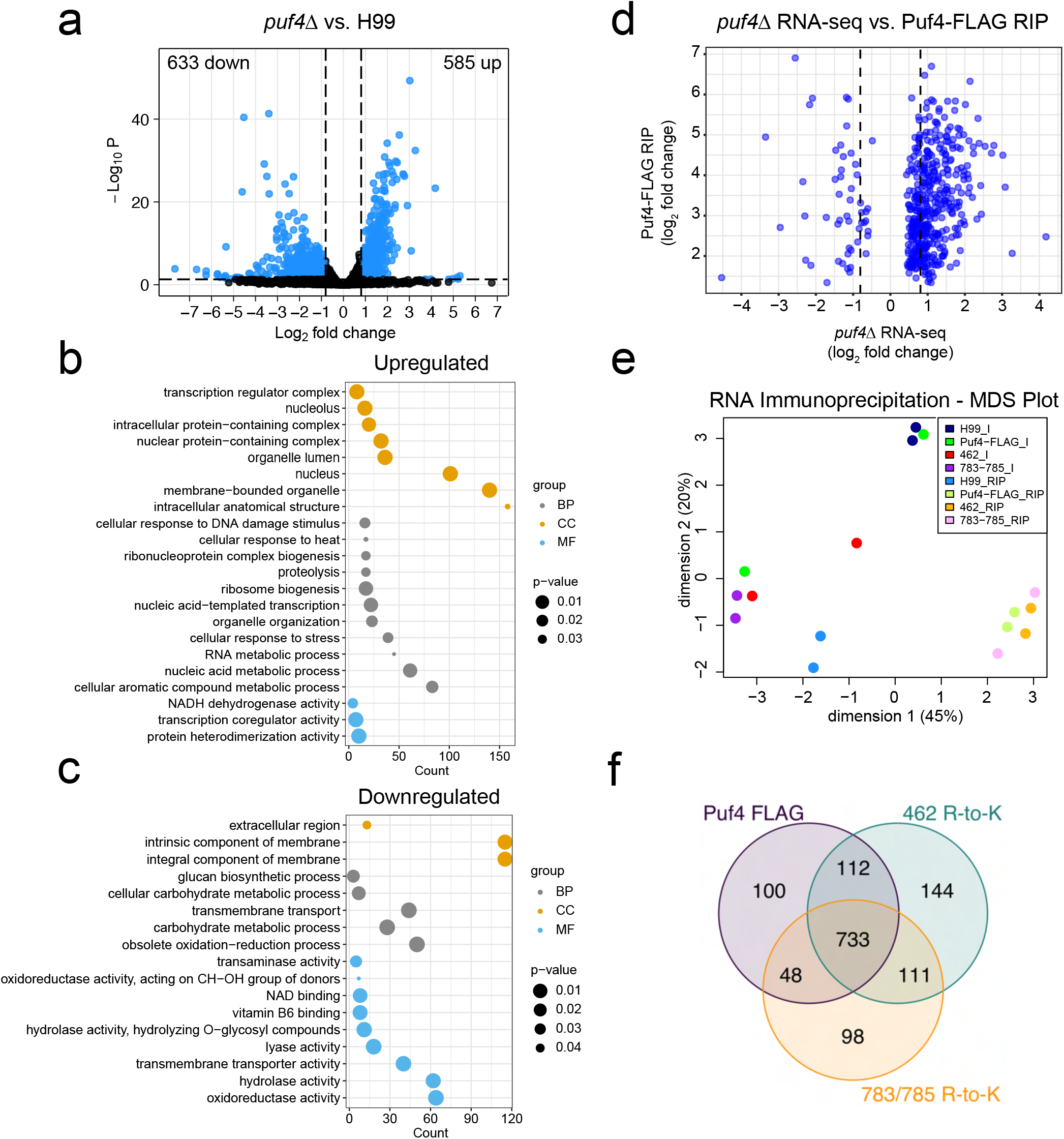
Diverse modulation of gene expression and RNA binding by Puf4 in *C. neoformans*. (a)Volcano plot summarizing the *puf4*Δ vs. wild type RNA-seq. Cells were grown to the mid-log stage in YPD broth at 30°C. Differentially expressed genes were identified using 1.75-fold difference and 0.05 adjusted *P* value thresholds. (b) Curated GO analysis of the upregulated genes. GO term analysis was performed using FungiDB. BP, biological process; CC, cellular component; MF, molecular function. (c) Curated GO analysis of downregulated genes. (d) Puf4 FLAG RIP vs. *puf4*Δ RNA-seq. Puf4-FLAG overexpression cells were grown overnight at 30°C to an optical density at 600 nm of 1.5–1.8. RNAs bound by Puf4 at 30°C were analyzed for their expression in the *puf4*Δ mutant to show that the majority of the Puf4-bound RNAs are negatively regulated by Puf4. (e) Multidimensional scaling (MDS) plot of RIP samples. (f) Venn diagram of RNAs bound by Puf4-FLAG, Puf4-FLAG 462 RtoK clone 1, and Puf4-FLAG 783/785 RtoK clone 2.

We then explored which of the transcripts are directly regulated by Puf4. We performed RNA immunoprecipitation (RIP) using the Puf4-FLAG overexpression strain to isolate the population of RNAs that are bound to Puf4 and the methyl-deficient mutant strains to define methylation-dependent interactions. Fold-enrichment of co-immunoprecipitated RNA was calculated relative to the input for each condition using 2-fold enrichment and 0.05 adjusted *P*value thresholds. We looked at the RNA-seq data from *puf4*Δ cells to determine how the Puf4-bound RNAs are regulated: most of the transcripts were upregulated in *puf4*Δ cells, suggesting that Puf4 functions primarily as a negative regulator of mRNA abundance (Figure 4d). This led us to speculate that the majority of the upregulated RNAs in the *puf4*Δ RNA-seq data were regulated through direct binding, whereas the downregulated genes were likely regulated indirectly via a transcription factor or another RBP.

In the multidimensional scaling graph, RIP samples clustered together (Figure 4e), suggesting that the populations of Puf4-bound RNAs are similar for the wild-type and methyl-deficient proteins. Of the 993 RNAs that were bound to wild-type Puf4-FLAG in the RIP assay, 733 also bound to the RtoK mutant proteins (Figure 4f), indicating that arginine methylation of Puf4 regulates a small proportion of RNA binding. A small number of the unique binding interactions were identified with high confidence as not only by a *P* value criteria but using a 2fold threshold of enrichment in only given strain and not others (Table 1; Figure S2). Interestingly, we found that only Puf4-FLAG 462 RtoK bound to ESCRT-II complex subunit Vps22 mRNAs and two vacuolar protein mRNAs. Because this mutant also did not complement tunicamycin resistance, methylation of this residue may determine whether Puf4 regulates mRNAs encoding proteins in the secretory pathway.

**Table 1.** Specific Puf4-RNA interactions

| Gene ID | Description |
| --- | --- |
| **Puf4-FLAG** |  |
| CNAG_12982 | lncRNA |
| CNAG_08020 | Hypothetical, *C. neoformans* specific |
| CNAG_04500 | Hypothetical, basidiomycete specific |
| CNAG_06924 | β-Fructofuranosidase (Suc2) |
| CNAG_04818 | Monocarboxylate transporter (Mch2) |
| CNAG_07360 | Proline rich protein, basidiomycete specific |
| CNAG_03976 | Basic-leucine zipper domain protein (Gsb1), basidiomycete specific |
| CNAG_02136 | Ribosomal protein s6 |
| CNAG_02956 | Transcription factor lws1 |
| CNAG_07746 | Methylenetetrahydrofolate dehydrogenase |
| CNAG_02877 | Transcription factor Fzc51, basidiomycete specific |
| CNAG_01577 | Glutamate dehydrogenase |
| CNAG_01027 | Succinate-semialdehyde dehydrogenase |
| **Puf4-FLAG 462 RtoK** |  |
| CNAG_12107 | lncRNA |
| CNAG_12620 | lncRNA |
| CNAG_05704 | ESCRT-II complex subunit VPS22 |
| CNAG_04878 | Transcription factor Fzc1, basidiomycete specific |
| CNAG_03250 | Duf202-domain containing vacuolar protein, basidiomycete specific |
| CNAG_07889 | Hypothetical protein |
| CNAG_02584 | Serine/threonine-protein phosphatase 2A activator 1 |
| CNAG_01491 | Ankyrin repeat-containing domain containing protein |
| CNAG_04141 | Vacuolar protein, carboxypeptidase |
| **Puf4-FLAG 783/785 RtoK** |  |
| CNAG_07922 | Transcription factor Fzc4, basidiomycete specific |
| CNAG_06667 | Hypothetical protein, basidiomycete specific |
| CNAG_06654 | Sigma intracellular receptor 2 domain containing protein |
| CNAG_01357 | Mitogen-activated protein kinase tyrosine protein phosphatase SDP101 |
| CNAG_07416 | Exosome complex component RRP41 (yeast Ski6) |
| CNAG_05553 | Transcription initiation factor TFIID component TAF4-domain containing protein, basidiomycete specific |
| CNAG_06134 | Basic-leucine zipper domain protein (Bzp1), basidiomycete specific |
| CNAG_00644 | Sphingolipid delta-4 desaturase |
| CNAG_00329 | NADH dehydrogenase (ubiquinone) 1 beta subcomplex 8 |
| CNAG_04136 | Methylthioribose-1-phosphate isomerase (Mri1) |
| CNAG_03142 | Hypothetical protein |

<sup>a</sup>Specific interactions after filtering for high confidence (as outlined in Figure S2).

### Puf4 deletion and methyl-deficient complement strains have a melanization defect associated with impaired induction of *LAC1* expression

A GO term analysis of all 993 Puf4-bound RNAs revealed which processes these RNAs are involved in, including nucleic acid metabolic processes and responses to ER stress induced by tunicamycin and temperature stress (Figure 5a). Additionally, gene regulation through histone modifications and RNA polymerase complex were also represented in the GO analysis. Interestingly, three of the biological processes identified are related to key virulence factors of *C. neoformans,* namely, response to temperature stimulus, negative regulation of capsule organization, and regulation of melanin biosynthetic process, which correspond to the thermotolerance, capsule production, and melanin production, respectively. We therefore investigated whether Puf4 deletion or methylation deficiency alters melanization in cells spotted on asparagine agar containing L-DOPA (3,4-dihydroxy-L-phenylalanine). The deletion mutant had an obvious deficiency in melanin production, and this defect was complemented by singlecopy Puf4-FLAG and Puf4-FLAG overexpression (Figure 5b). However, the methyl-deficient mutants did not complement the melanization defect.

**Figure 5.**
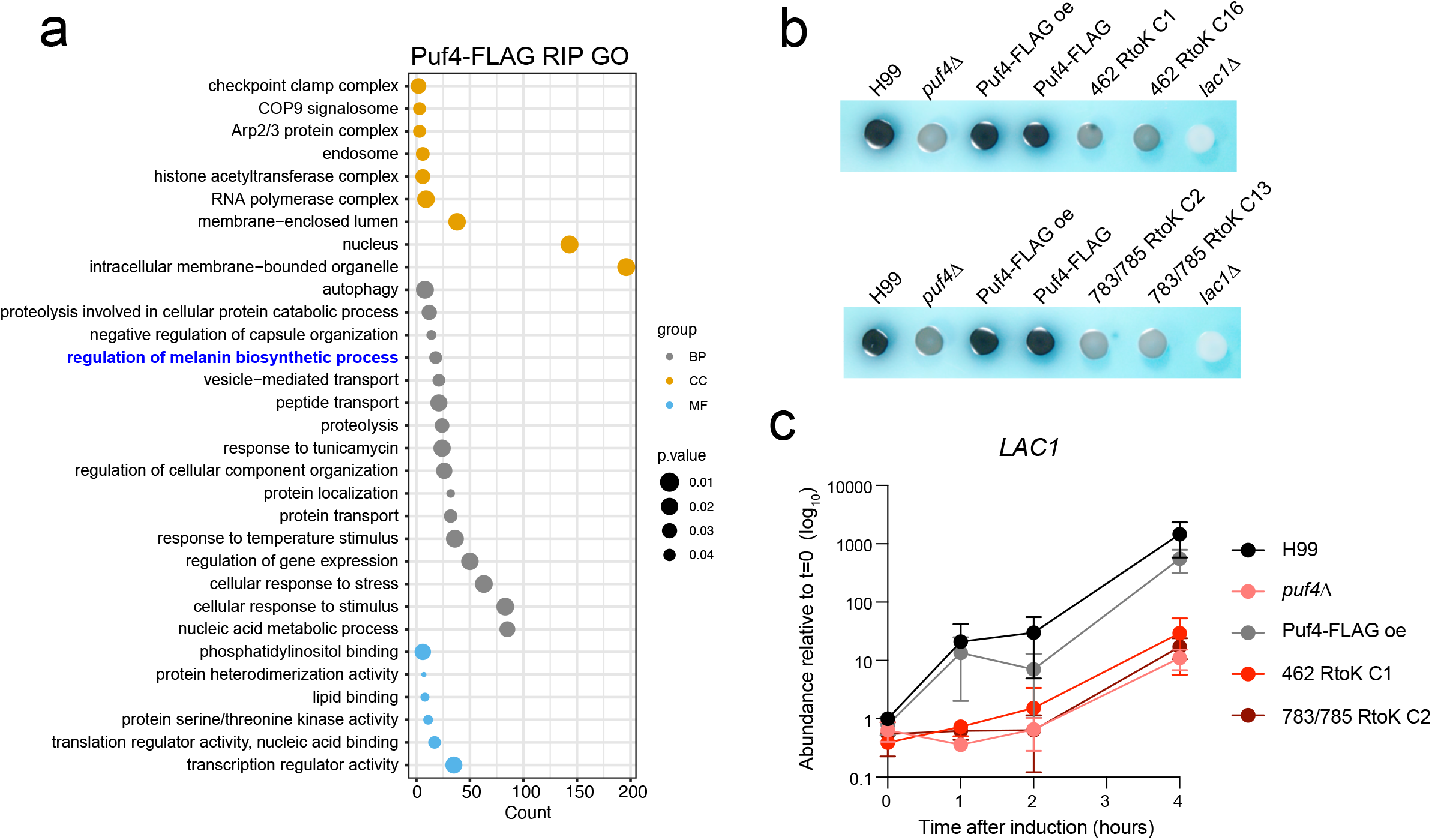
Puf4 deletion and methyl-deficient complement strains have a melanization defect associated with impaired induction of *LAC1* expression. (a) GO term analysis of RNAs bound by Puf4-FLAG. BP, biological process; CC, cellular component; MF, molecular function. (b) Melanin production. Cells were incubated overnight on asparagine agar supplemented with l-DOPA. The *lac1*Δ mutant was included as a negative control. (c) *LAC1* induction. Cells were grown to mid-log phase in YPD broth and transferred to asparagine-containing medium (for carbon starvation) for 4 h. *LAC1* abundance was determined by RT-qPCR at the indicated time points and *GPD1* was used as the normalization gene.

We next examined the strains for expression of *LAC1,* which encodes the multi-copper oxidase laccase that catalyzes the oxidation of L-DOPA to produce melanin ^38^. LAC1 expression is induced by glucose withdrawal; thus, we performed a glucose starvation time course experiment over 4 h. The wild-type strain and the Puf4-FLAG overexpression complement strain showed robust induction of *LAC1,* whereas the *puf4*Δ and methyl-deficient mutants did not (Figure 5c). This suggests that the inability to induce *LAC1* expression underlies the observed melanization defect.

### Methyl-deficient mutants have altered protein-protein interactions and PTM landscape

To further evaluate the role of Puf4 in orchestrating various post-transcriptional events, we investigated the protein interaction networks of Puf4 at 30°C and 37°C. Because the *puf4*Δ mutant has a defect in thermotolerance, we reasoned that we would be able to discern temperature-specific protein interaction networks. We immunoprecipitated Puf4 from the Puf4-FLAG overexpression strain and the methyl-deficient strains (Puf4-FLAG 462 RtoK clone 1 and Puf4-FLAG 783/785 RtoK clone 2) and identified interacting proteins via MS. We used stringent criteria for our MS data filtering and only considered proteins with at least three spectral counts and 2-fold enrichment over that in an untagged mock immunoprecipitation. Additionally, interactions that were captured in only a single biological replicate were excluded. Our results revealed that Puf4 has temperature-specific protein-protein interactions (Figure 6a and 6b). Puf4 interacted with Ago1 (argonaute-1), Upf1 (nonsense-mediated decay helicase), eIF4A (translation initiation factor), Not1 (Ccr4-Not complex subunit), Pab1 [poly(A) binding protein], Cid1 [poly(A) polymerase], Bmh2 (14-3-3 domain-containing protein), RNaseH1, and Rpt1 (proteasome subunit) at 30°C (Figure 6c). Puf4 interacted with additional sets of proteins at 37°C, including Lin1 (CNAG_O4368; U5 snRNP complex protein), Iws1 (transcription factor), Kap123 (importin), Hrp1 (pre-mRNA cleavage and polyadenylation-related protein), Dbp2 (pre-mRNA splicing protein), and Hsp90 (heat shock chaperone) (Figure 6d). These results suggest a that Puf4 interacts with proteins involved in cytoplasmic processes (such as Not1) at 30°C but begins to interact with proteins involved in nuclear RNA processes at higher temperatures.

**Figure 6.**
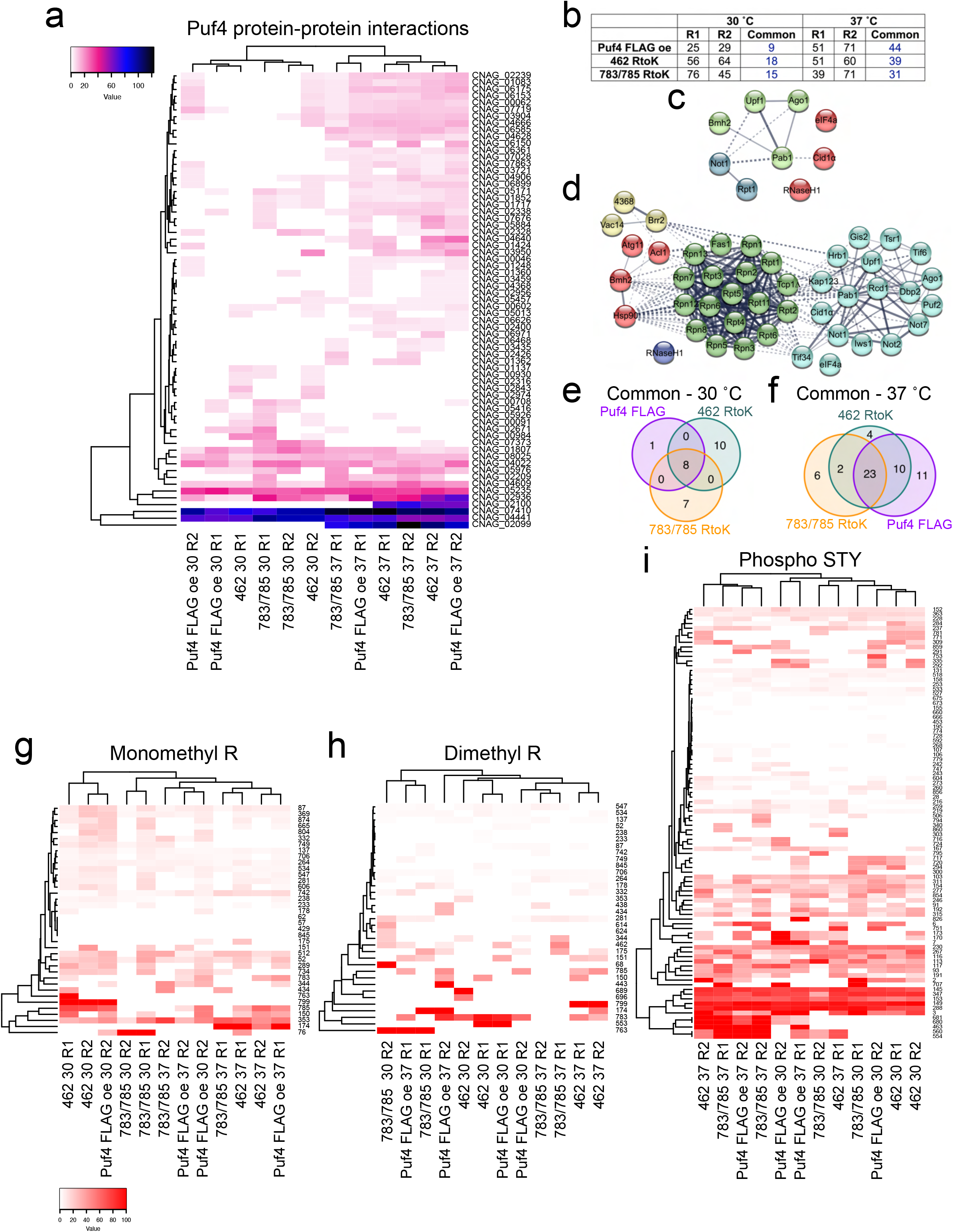
Methyl-deficient mutants have altered protein–protein interactions and PTM landscape. (a) Heat map of spectral counts from IP/MS. Puf4-FLAG overexpression strain and methyl-deficient mutants were grown to mid-log phase at 30°C or 37°C in YPD broth. FLAG immunoprecipitation was performed to capture protein-protein interactions, and interacting proteins were identified using LC-MS/MS in two biological replicates. Untagged cells were used to detect nonspecific interactions. (b) Summary table. Hits with at least three spectral counts 2-fold over the background amount were selected for each replicate. Common column shows the number of proteins that were common between two replicates. (c,d) Protein-protein interaction maps of 9 proteins that interacted with Puf4-FLAG at 30°C and 44 proteins that interacted with Puf4-FLAG at 37°C. (e,f) Venn diagrams of common interactors in both replicates of Puf4-FLAG and methyl-deficient mutants at 30°C and 37°C. (g-i) PTM landscape. Monomethyl arginine, dimethyl arginine, and STY phosphorylation of Puf4 were identified in Puf4-FLAG or methyl-deficient mutants to identify cross talk between PTMs.

We identified 26S protease regulatory subunit 7 (*S. cerevisiae* Rpt1) as the single protein unique to the interactome of Puf4-FLAG (Figure 6e). *S. cerevisiae* Rpt1 is an essential gene and is a multifunctional protein involved in protein catabolism as well as RNA polymerase II preinitiation complex assembly. Ten proteins were unique to the interactome of Puf4-FLAG 462 RtoK at 30°C these included mitochondrial proteins and a cytoplasmic protein involved in glucose metabolism. Puf4-FLAG 783/785 RtoK had seven unique interactions, but four were included in one replicate of Puf4-FLAG at 30°C. The three interactions that we can report with high confidence included Cpa2 (carbamoyl-phosphate synthase) and Rcd1 (cell differentiation protein). At 37°C, four proteins were found to interact with Puf4-FLAG 462 RtoK (Figure 6f), but only one of these was not present in a replicate of a difference strain: myosin class V heavy chain. The five unique high-confidence interactions with Puf4-FLAG 783/785 RtoK at 37°C were nuclear proteins. These findings suggest that Puf4 has temperature- and methylation-dependent protein interaction networks.

We were able to identify the PTMs of the Puf4 immunoprecipitated from the methyl-deficient mutants. RtoK mutation(s) at either Arg462 or Arg783/Arg785 impacted the methylation of other residues (Figure 6g and 6h). For example, Arg799 was more strongly monomethylated in the Puf4-FLAG 462 RtoK mutant at 30°C than in the wild-type or 783/785 RtoK mutant Puf4 proteins and was strongly dimethylated in the Puf4-FLAG 462 RtoK mutant at 37°C. This suggests that mutation (or modification) of a critical arginine residue (e.g., Arg462) impacts the modification of other arginines.

We also examined whether other PTMs are affected in our methyl-deficient mutants. Specifically, we looked at changes in STY phosphorylation (see Figure 1d), because the cross talk between methylation and phosphorylation may influence protein interactions. Our results showed that methyl-deficient mutations alter the phosphorylation of multiple domains. For example, Thr93 was not phosphorylated in the Puf4-FLAG 783/785 RtoK mutant. This residue is located C terminal to the coiled-coil domain in a disordered domain that may influence phase separation of this RBP. The temperature-dependent phosphorylation of some residues (e.g. Thr554 and Ser560 phosphorylated at 37°C but not at 30°C) in the Puf4-FLAG overexpression cell line may also influence the protein interactions at different temperatures.

## Discussion

Although the PUF domains in RBPs are evolutionarily conserved, we found that the RG-rich domains, which may serve as methylation hubs, are not. Mutations that interfered with arginine methylation at specific residues in Puf4 did not grossly impact RNA binding but did affect mRNAs important for antifungal resistance, thermotolerance, and ER stress. Our data suggest that the PTMs in the RG-rich domain of Puf4 fine-tune the protein-protein interactions that regulate these processes ^26,27^

The identification of methylation-dependent roles for Puf4 in cellular processes important for *C. neoformans* reveals PTM-driven unique functional states. For example, we observed differential complementation of some *puf4*Δ phenotypes by the two methyl-deficient mutants. The concept of PTM-driven functional diversity may explain why deletion of a gene that regulates essential cellular processes such as ribosome biogenesis and translation does not always produce a drastic phenotype. Multiple subtly different PTM-driven functional states may neutralize or balance each other, effects that are masked by full deletion of the target. Further biochemical and cellular assays using various arginine mutants and recombinant proteins are necessary to further explore this hypothesis.

Our complementation experiments with methyl-deficient Puf4 mutants indicated gene dose-dependent effects on the antifungal resistance, mRNA stability, and cell wall remodeling phenotypes observed in *C. neoformans*. This suggests to us that the changes in functional states can be rather subtle, and increased expression can skew the effect on downstream targets. By contrast, the inability of the methyl-deficient mutants to complement the thermotolerance defect of the *puf4*Δ strain at 38°C suggests that the modification of arginines in Puf4 drives temperature-specific protein function. This is especially intriguing because *RMT5*(which interacts with Puf4) is the only *C. neoformans* arginine methyltransferase for which deletion similarly eliminates thermotolerance at elevated temperatures ^25^. Thus, we speculate that Rmt5 acts on Puf4 to establish thermotolerance.

Our previous work showed that increased cell wall chitin content is one of the factors driving caspofungin resistance in *C. neoformans*^26^. Surprisingly, we discovered that overexpression of Puf4 increased cell wall chitin and chitooligomers similarly to *PUF4* deletion. However, the strain overexpressing Puf4 was not resistant, and even modestly hypersensitive, to caspofungin. This highlights the importance of gene dosage and suggests that overexpression of Puf4 may drive a dominant negative function or interactions with other regulatory molecules.

Our transcriptomic analysis revealed that Puf4 regulates the expression of many genes; in general, genes related to gene regulation were upregulated and genes involved in carbohydrate metabolism and membrane components were downregulated. Notably, most of the differentially expressed lncRNAs in the *puf4*Δ strain were downregulated. In humans, lncRNAs act as endogenous competitors to prevent Pumilio domain-containing proteins from binding to their target mRNAs ^39,40^. This regulatory mechanism may therefore be evolutionarily conserved. By contrast, Puf4 and other Pumilio domain-containing proteins also exhibit evolutionary diversity, such as in fungal clade-specific “rewiring” of the Puf4 post-transcriptional regulon ^14,41^. For example, Puf4 regulates nucleolar RNAs in the clade *Saccharomycotina* but mitochondrial RNAs in the clade *Pezizomycotina*.

The GO analysis indicated that Puf4 is important for secretory processes in *C. neoformans*. Accordingly, the *puf4*Δ and methyl-deficient mutant strains had defects in melanization (melanin is a secreted factor). This defect was accompanied by disrupted regulation of *LAC1* expression. Because *LAC1* mRNA contains Puf4 binding elements in its 5′ leader and 3′ trailer regions, we posited that *LAC1* mRNA was targeted by Puf4 under the experimental conditions (i.e., carbon starvation); the absence of *LAC1* in our RIP-seq data is consistent with this. The magnitude of *LAC1* induction may indicate that Puf4 broadly regulates transcription of *LAC1* through histone modifications, RNA polymerase complex, and/or transcription factors. For example, the transcription factor Hob1 controls melanization in *C. neoformans,* and we observed *HOB1* mRNA bound to Puf4 in our RIP-seq analysis ^42^. Our previous work showed that post-transcriptional events drive transcriptome remodeling, and a defect in transcription factor translation impacts subsequent gene expression ^34^. Future work on Puf4-regulated transcription factors will shed more light on post-transcriptional regulation virulence factor production in this important human pathogen.

Our RIP-seq analysis also identified transcripts that were specifically bound by wild-type Puf4 and methyl-deficient Puf4 mutants. Even though these were small populations of specific binding interactions, changes in interactions of one target may be responsible for an observed phenotype. Therefore, we plan to explore these specific interactions in the future to delineate their biological impacts.

Our analyses of protein networks revealed temperature-dependent protein-protein interactions with the wild-type Puf4 and its methyl-deficient mutants. Of note we found that Puf4 interacts with Ccr4/Not and proteasome complexes, which suggests that it is a central regulator of mRNAs marked for degradation, perhaps as a mitigator of cellular stress or damage. Our analyses also revealed putative methylation-dependent interactions. Some of these interactions may be mediated by PTM cross talk, as our results suggest there is cross talk involving different methylation sites and between sites of methylation and phosphorylation. Indeed, PTM cross talk is an important aspect of functional regulation and diversity ^16,43–45^. One example of PTM crosstalk is the dynamic regulation of chromatin states through histone H3 acetylation and phosphorylation ^47^.

Although our approach to characterize the PTM landscape of Puf4 yielded high protein coverage and we were confidently able to identify methyl modifications to arginines, we were unable to distinguish between symmetric and asymmetric dimethylation, which can determine how the residue interacts with a “reader” of PTMs. Future studies will include MS approaches that can more confidently detect these differences (reviewed 41). The use of RtoK mutations to dissect the functional importance of modified arginine residues enabled us to conclude that the presence of lysine instead of arginine changes protein function, but we cannot definitely state that the effect is the result of a lack of arginine methylation. Studies on the effects of mutating those arginines to phenylalanines to mimic methylation or to alanines would validate the importance of the modified residues.

The outlined studies and findings in this article explore the post-transcriptional regulation network of Puf4 comprehensively through a specific lens of protein arginine methylation, evolution, and fungal virulence factors. These results will serve as the basis of future investigations on PTM-driven post-transcriptional events in *C. neoformans*.

## Acknowledgments

We acknowledge Dr. Amanda L. M. Bloom for influential discussions and Dr. Runpu Chen for advice on sequencing data analysis. We thank Dr. Karen Dietz for editing and proofreading. This work was funded, in part, by American Heart Association (grant no. 33440094 and predoctoral fellowship 827110 [to M.C.K.]) and the National Institute of Allergy and Infectious Disease (grant no. R21AI33133 to J.C.P.).

## Author contributions

Conceptualization, M.C.K. and J.C.P.; Investigation, M.C.K., S.D. and S.S.; Resources, J.N.K., J.Q.; Writing–Original Draft, M.C.K.; Writing–Review& Editing, M.C.K. and J.C.P.; Visualization, M.C.K.; Supervision, J.C.P.; Funding Acquisition, J.C.P.

## STAR Methods

### EXPERIMENTAL MODEL AND SUBJECT DETAILS

#### Yeast strains and molecular cloning

Strains used in this study were derived from *Cryptococcus neoformans* var. *grubii* strain H99, a fully virulent strain (gifted by Peter Williamson, UIC, NIAID) derived from strain H99O (gifted by John Perfect, Duke University). The *puf4*Δ mutant and Puf4-FLAG complement cell lines were previously established ^26^. Methyl-deficient Puf4 constructs were established using a QuickChange site-directed mutagenesis kit (Agilent). All primers used in this study are listed in Table S1. Cell lines were established using biolistic transformation.

### METHOD DETAILS

#### Phylogenetic analysis

A BLAST search was used to find the orthologs of Rmt5 in fungal species from *Ascomycota, Basidiomycota, Mucoromycotina,* and *Chytridiomycota* phyla by using the *C. neoformans PUF4* gene (CNAG_02810) as the query. All sequences were aligned using the default algorithm of Geneious alignment followed by a neighbor-joining tree protein alignment on Geneious. Human *PUM1* and *PUM2* were included as outgroups.

#### Puf4 immunoprecipitation and LC-MS/MS

##### For PTM analysis

FLAG immunoprecipitation was performed as described previously ^26^. Puf4-FLAG overexpression strain and wildtype (H99) cells were grown to an optical density at 600 nm (OD_600_) of 1.5–1.8 at 30°C in 2 liters YPD broth while shaking at 200 rpm. Cells were collected by centrifugation at 7,000 ×*g* for 5 min. Pellets were frozen in liquid N2 and stored at −80°C. Frozen cell pellets were ground using liquid N2 in a coffee grinder (Krups F203) for 3 min. Cell powder was then further ground using a mortar and pestle for 15–20 min with the frequent addition of liquid N2. Cell powder was transferred to 50-ml conical tubes and stored at −80°C or immediately used for immunoprecipitation. Cell powder was dissolved in RIP buffer (25 mM HEPES-KOH [pH 7.9], 0.1 mM EDTA, 0.5 mM EGTA, 2 mM MgCl_2_, 20% glycerol, 0.1% Tween-20, 300 mM KCl, 1×cOmplete protease inhibitor [Roche], 50 U/ml RNaseOUT [Invitrogen]) and incubated on a rotator for 1 h at 4°C. The lysate was cleared by centrifuging at 27,000 ×*g* for 20 min at 4°C. Cleared lysate was incubated with anti-FLAG antibody-coated magnetic agarose beads (Pierce) for 4 h at 4°C. The beads were then immobilized by magnet and washed four times with RIP buffer. Protein was eluted twice using 50 μl of elution buffer (0.1 M glycine [pH 2]). Elution volumes were separated on 4–15% Mini-PROTEAN TGX stain-free precast polyacrylamide gels (Bio-Rad) at 150 V. Gels were stained with Coomassie blue, and bands corresponding to Puf4-FLAG were excised. Excised bands were stored at −80°C until trypsin digestion and LC-MS/MS analysis.

##### Forprotein–protein interactions

Immunoprecipitations using the Puf4-FLAG overexpression strain, Puf4-FLAG 462 RtoK clone 1, Puf4-FLAG 783/785 RtoK clone 2, and H99 cells were performed as described above. Instead of glycine elution, bound proteins were eluted using FLAG elution buffer (25 mM HEPES-KOH [pH 7.9], 2 mM MgCl_2_, 20% glycerol, 300 mM KCl, 1×cOmplete protease inhibitor (Roche), 1×PhosSTOP phosphatase inhibitor, and 0.5 mg/ml 3×FLAG peptide). After washing four times with RIP buffer (15-min each), the beads were eluted three times using 100 μl of FLAG elution buffer. Elution volumes were combined and stored at −80°C until trypsin digest and LC-MS/MS analysis. Prior to LC-MS/MS analysis, the buffer was exchanged for one compatible with trypsin digest and without glycerol.

##### Protein digestion

A surfactant-aided precipitation/on-pellet digestion protocol was adopted using our previously published method with slight modification ^48^. Samples were transferred to 10-kDa MWCO centrifugal filter units (MilliporeSigma) for buffer exchange and concentration. Three consecutive centrifugation steps with the addition of 200, 400, and 400 μl 50 mM Tris-formic acid (FA) (pH 8.4) were performed at 14,000 ×*g* at room temperature for 20 min, and samples were concentrated to a final volume of 20 μl. A total of 60 μl 0.5% SDS was then added to each sample, and the filter units were vortexed vigorously for 10 min. Concentrated samples were transferred to LoBind microcentrifuge tubes (Eppendorf). Proteins were sequentially reduced by 10 mM dithiothreitol at 56 °C for 30 min and alkylated by 25 mM iodoacetamide at 37°C in darkness for 30 min. Both steps were performed in a thermomixer (Eppendorf) with rigorous shaking. Proteins were then precipitated by adding 6 volumes of chilled acetone with vortexing, and the mixture was incubated at −20°C for 3 h. Samples were then centrifuged at 20,000 ×*g* at 4°C for 30 min, and the supernatant was removed. Protein pellets were gently rinsed with 500 μl methanol, air-dried for 1 min, and resuspended in 46 μl 50 mM Tris-FA (pH 8.4). A total volume of 4 μl trypsin (Sigma Aldrich), reconstituted in 50 mM Tris-FA (pH 8.4) to a final concentration of 0.25 μg/μl, was added for a 6-h tryptic digestion at 37°C with constant shaking in a thermomixer. Digestion was terminated by adding 0.5 μl FA, and samples were centrifuged at 20,000 ×*g* at 4°C for 30 min. The supernatants were carefully transferred to LC vials for analysis.

##### LC-MS analysis

The LC-MS system consists of a Dionex UltiMate 3000 nano-LC system, a Dinoex UltiMate 3000 micro-LC system with a WPS-3000 autosampler, and an Orbitrap Fusion Lumos mass spectrometer. A large-inner diameter (i.d.) trapping column (300-μm i.d.×5 mm) was implemented before the separation column (75-μm i.d. ×65 cm, packed with 2.5-μm XSelect CSH C 18 material) for high-capacity sample loading, cleanup, and delivery. For each sample, 12 μl derived peptides was injected twice consecutively for LC-MS analysis. Mobile phase A and B were 0.1% FA in 2% acetonitrile and 0.1% FA in 88% acetonitrile, respectively. The 180-min LC gradient profile was 4% phase B for 3 min, 4%-9% phase B for 5 min, 9%-31% phase B for 117 min, 31%-50% phase B for 10 min, 50%-97% phase B for 1 min, 97% phase B for 17 min, and then equilibrated to 4% phase B for 27 min. The mass spectrometer was operated under data-dependent acquisition mode with a maximal duty cycle of 3 s. MS1 spectra were acquired by Orbitrap under 120k resolution for ions within the *m/z* range of 400-1,500. automatic gain control and maximal injection time were set at 175% and 50 ms, respectively, and dynamic exclusion was set at 60 ± 10 ppm. Precursor ions were isolated by quadrupole using a *m/z* window of 1.6 Th and were fragmented by high-energy collision dissociation. MS2 spectra of a precursor ion fragmented were acquired by a back-to-back Orbitrap and Ion Trap scheme. Orbitrap was operated under 15k resolution with an automatic gain target target of 100% and a maximal injection time of 35 ms; Ion Trap was operated under rapid scan rate with an automatic gain control target of 100% and a maximal injection time of 50 ms. Detailed LC-MS settings and relevant information can be found in a previous publication by Shen et al. ^49^.

##### Data processing

To identify the Puf4 interactome and characterize PTMs of Puf4, two separate sets of data processing procedures were applied to the LC-MS files. For interactome identification, LC-MS files were searched against *Cryptococcus neoformans* var*. grubii* serotype A Swiss-Prot+TrEMBL protein sequence database (7,429 entries) using Sequest HT embedded in Proteome Discoverer 1.4 (Thermo Fisher Scientific). A target-decoy approach using a concatenated forward and reverse protein sequence database was applied for false-discovery rate estimation and control. The searching parameters included (i) precursor ion mass tolerance, 20 ppm; (ii) product ion mass tolerance, 0.8 Da; (iii) maximal missed cleavages per peptide, 2; (iv) fixed modifications, cysteine carbamidomethylation; (v) dynamic modifications, methionine oxidation and peptide N-terminal acetylation. Search result merging, protein inference/grouping, and false-discovery rate control were performed in Scaffold 5 (Proteome Software, Inc.). For identification, the global protein/peptide false-discovery rate was set to 1.0% and at least two unique peptides were required for each protein. For quantification, protein abundance was determined by total spectrum counts and total MS2 ion intensities. Results were exported and manually curated in Microsoft Excel.

For Puf4 PTM characterization, LC-MS files were searched against *Cryptococcus neoformans* Puf4 sequence (UniProt protein accession J9VI91) using the same search engine settings with the following additional parameters. For Puf4 methylation, arginine (R) methylation and dimethylation were included in dynamic modifications; for Puf4 phosphorylation, serine (S)/threonine (T)/tyrosine (Y) phosphorylation was included in dynamic modifications. Search results were filtered to only retain peptides with medium and high confidence and were exported from Proteome Discoverer. Data curation was performed using a customized R script.

#### Spot plate dilution assay

Cells were grown overnight at 30°C in YPD broth. Overnight cultures were washed with sterile distilled water, and the OD6°°was adjusted to 1 in water. Adjusted cultures were 1:10 serially diluted five times, and 5 μl of each dilution was spotted onto YPD agar plates containing the selected drugs. Agar plates were incubated for 2-3 days at the temperatures indicated in the text and photographed.

#### RNA stability time course

RNA extraction and stability analysis were performed as described previously ^26^. Briefly, midlog-stage cultures were supplemented with 250 mg/ml of the transcriptional inhibitor 1,10-phenanthroline (Sigma). Then, 5-ml aliquots of each culture were transferred to snap-cap tubes and pelleted every 15 min for 60 min. Fifty microliters RLT buffer supplemented with 1%ß-mercaptoethanol was added to each pellet prior to flash freezing in liquid N2. Pellets were stored at −80°C until RNA extraction. Cells were lysed by bead beating using glass beads. RNA was extracted from each sample using the RNeasy mini kit (Qiagen) according to the manufacturer’s instructions. RNA was DNase digested on column using the RNase-free DNase kit (Qiagen) or using the Ambion Turbo DNA-free kit (Thermo Fisher Scientific). cDNA for real-time quantitative PCR (RT-qPCR) was synthesized using the Applied Biosystems high-capacity cDNA reverse transcription kit (Thermo Fisher Scientific). Samples were quantified using the second-derivative maximum method and fitted to a standard curve of five 4-fold serial dilutions of cDNA. Primer sequences were previously published ^28^.

#### Cell wall composition flow cytometry

Cell wall chitin and exposed chitooligomer levels were quantified using flow cytometry after cells were stained with calcofluor and wheat germ agglutinin, as described previously ^26^. Mean fluorescence intensities were used to calculate the abundance of cell wall components relative to that in the wild type.

#### Melanin assay

Melanization agar plates were prepared using asparagine medium as described above and 2% agar supplemented with 2 mM 3,4-dihydroxy-L-phenylalanine (L-DOPA) at pH 6.5. Cells were grown overnight at 30°C, and OD600 was adjusted to 1 in water. Five microliters of each culture was spotted onto melanin agar plates, incubated overnight at 30°C, and photographed.

#### RNA-seq

RNA was extracted from the wild type and the *puf4*Δ mutant grown to mid-log phase at 30°C.

RNA extractions were performed as described above. RNA quality was determined by investigating integrity on an RNA gel and using a bioanalyzer prior to sequencing. RNA sequencing was performed as described previously ^25^. RNA samples were submitted to Genewiz for library preparation and Illumina sequencing. Two biological replicates were analyzed per strain. RNA sequencing libraries were prepared using the NEBNext Ultra II RNA library prep kit for Illumina according to the manufacturer’s instructions (NEB). RNA sequencing reads were trimmed to remove adapter sequences and filtered using Cutadapt. Filtered reads were aligned to the *C. neoformans* H99 genome downloaded from FungiDB using STAR alignment. After mapping, there were approximately 12–14 million reads per sample. The read counts for each gene were calculated using RSEM. Differentially expressed genes between the mutant and wild-type samples were determined using DeSeq2 in R. Data were then filtered according to a 1.75-fold change and an adjusted *P* value threshold of 0.05.

#### RIP-seq

Wild-type (H99), Puf4-FLAG overexpression, Puf4-FLAG 462 RtoK clone 1, and Puf4-FLAG 783/785 RtoK clone 2 cell lines were grown to an OD600 of 1.5–1.8 in 2-liter YPD cultures. FLAG immunoprecipitation was performed as described above. Immunoprecipitated proteins and bound RNAs were eluted using low-pH glycine elution buffer, and RNA was extracted from elution volumes using low-pH phenol-chloroform. Untagged wild-type cells were used as a negative control for nonspecific enrichment of RNAs. Additionally, RNA was extracted from input samples to utilize as a baseline for enrichment calculation. RNA samples were submitted to Genewiz for library preparation and sequencing. Two biological replicates per strain were analyzed. Reads were trimmed, filtered, and mapped as described above. The read counts for each gene were calculated using RSEM. Differentially expressed genes between the RIP and input samples were determined using DeSeq2 in R. A 2-fold enrichment and 0.05 adjusted *P*value threshold was used in these analyses. Any RNAs that were identified as enriched in the untagged control were excluded from analyses of all other samples because they were considered nonspecific interactions.

## QUANTIFICATION AND STATISTICAL ANALYSIS

For RNA stability analyses, statistical differences of decay rates were compared by determining the least-squares fit of one-phase exponential decay nonlinear regression analysis with GraphPad Prism software. Significance between curves was detected with a *P* value cutoff of 0.05, which determined that the data from two different curves create different regression lines, therefore yielding different half-lives of the same transcript investigated in different mutants.

## RESOURCE AVAILABILITY

### Lead contact

Further information and requests for resources and reagents should be directed to and will be fulfilled by the lead contact, John C. Panepinto.

#### Materials availability

All constructs and strains are available upon request.

#### Data availability

All RNA-seq and RIP-seq files for this project will be available in the NCBI Gene Expression Omnibus (GEO), accession number pending. Data tables for MS, RNA-seq and RIP-seq experiments are also included in the Table S1.

---

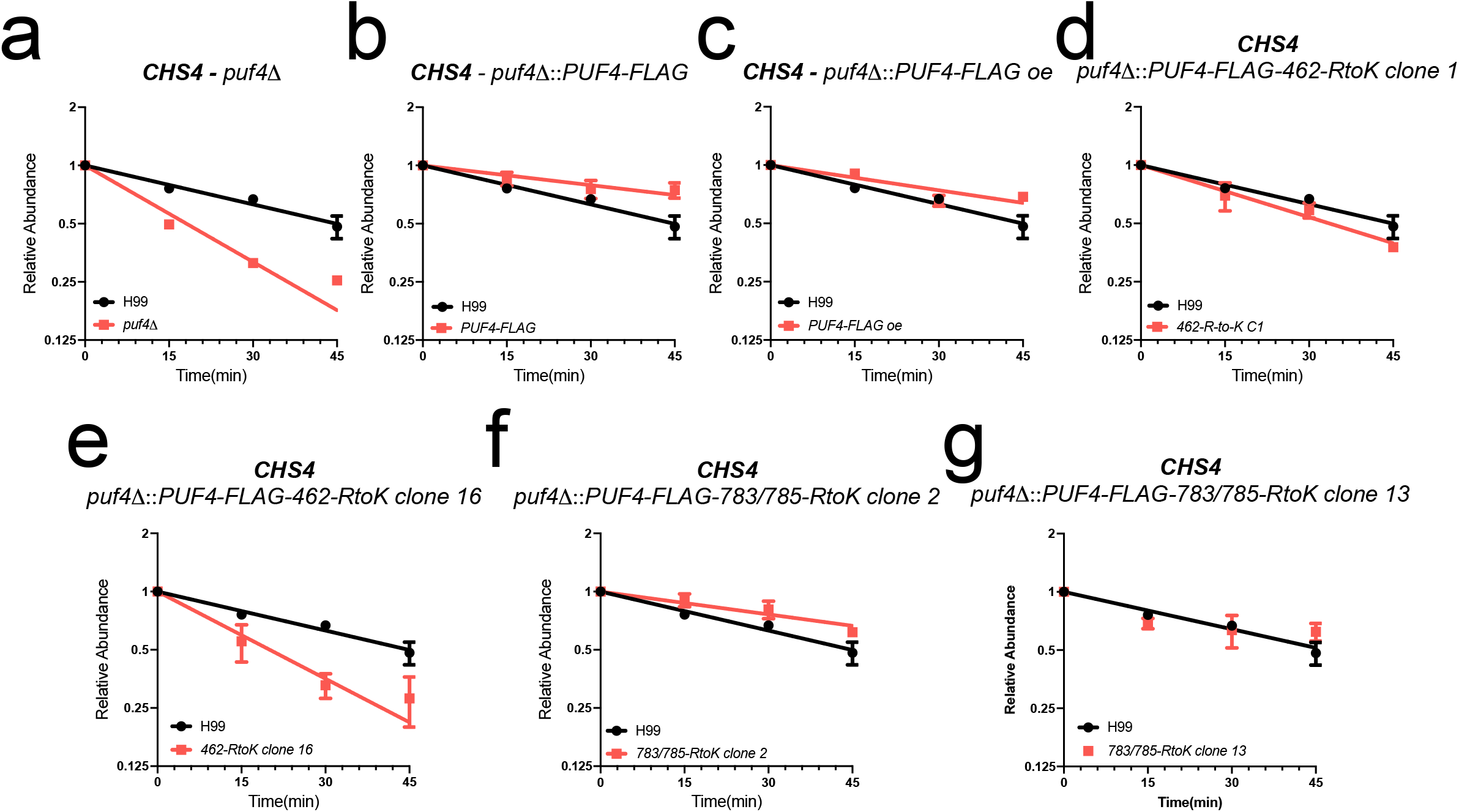

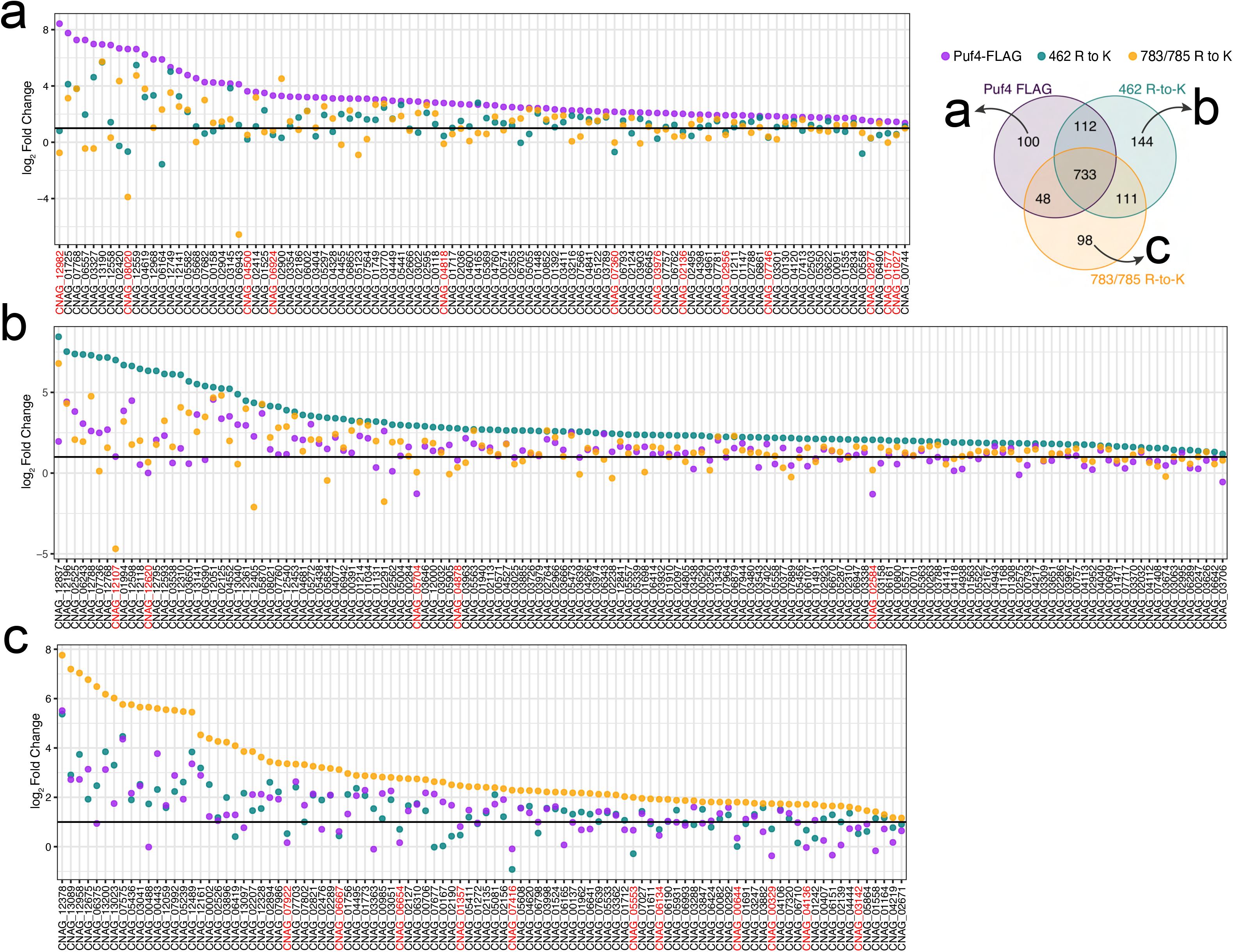

